# Ecological Observations Based on Functional Gene Sequencing Are Sensitive to the Amplicon Processing Method

**DOI:** 10.1101/2022.02.10.480020

**Authors:** Fabien Cholet, Agata Lisik, Hélène Agogué, Umer Z Ijaz, Philippe Pineau, Nicolas Lachaussée, Cindy J Smith

## Abstract

Until recently, the de-facto method for short read-based amplicons reconstruction is a sequence similarity threshold approach (Operational taxonomic Units OTUs). This assumption was relaxed by shifting to Amplicon Sequencing Variants (ASVs) where distributions are fitted to abundance profiles of individual genes using a noise-error model. Whilst OTUs-based approach is still useful for *16SrRNA/18S rRNA* regions, where typically 97-99% thresholds are used, their utility to functional genes is still debatable as there is no consensus on how to cluster the sequences together. Here, we compare OTUs- and ASVs-based reconstruction approaches as well as taxonomy assignment methods, Naïve Bayesian Classifier (NBC) and Bayesian Lowest Common Ancestor Algorithm (BLCA), using functional genes dataset from the microbial nitrogen-cycling community in the Brouage mudflat (France). A range of OTU similarity thresholds and ASV were used to compare *amoA* (AOA and AOB), *nxrB*, *nirS*, *nirK* and *nrfA* communities between differing sedimentary structures. We show that for AOA-*amoA* and *nrfA*, the use of ASV led to differences in the communities between sedimentary structures whereas the use of OTUs didn’t. Conversely, significant differences were detected when using OTU (97%) for AOB-*amoA* but not with ASV or OTUs at other similarity thresholds. Interestingly, conclusions drawn from the other three functional genes were consistent between amplicon reconstruction methods. We also show that, when the sequences in the reference-database are related to the environment in question, BLCA leads to more phylogenetically relevant classifications. However, when the reference database contains sequences more dissimilar to the ones retrieved, NBC helps obtain more information.

**Importance:** Several analysis pipelines are available to microbial ecologists to process amplicon sequencing data yet to-date, there is no consensus as to the most appropriate method, and it becomes more difficult for genes that encode a specific function (functional genes). Standardised approaches need to be adopted to increase reliability and reproducibility of environmental amplicon sequencing-based datasets. In this paper, we argue that the recently developed ASV approach offers a better opportunity to achieve such standardisation compared to OTUs for functional genes. We also propose a comprehensive framework for quality filtering of the sequencing reads based on protein sequence verification and merging.

## Introduction

The Polymerase Chain Reaction (PCR) combined with High Throughput Sequencing (HTS) has revolutionised our understanding of microbial ecology (1). Amplicon sequencing of the *16S rRNA* gene as a molecular marker for diversity is now routine (2). The approach can also be applied to genes and/or transcripts encoding enzymes for specific functions (functional genes). Functional genes involved in cell division and maintenance (housekeeping genes) can be used as an alternative to the *16S rRNA* for diversity estimation (3, 4) to circumvent the issue of intra-genomic variation in the *16S rRNA* gene (5). Biogeochemical cycles can also be targeted via key functional genes involved in the processes of interest *e.g.* the nitrogen cycle (6–11); sulphur cycle (12, 13); methane cycle (14). The same is true for bioremediation (15, 16) and antibiotic resistance (17) to mention but a few possibilities. By targeting functional genes we can start to unravel the functional potential of microbial communities, and if transcripts are targeted, actively transcribing organisms are revealed, a step closer to identifying the organisms driving the target processes.

Irrespective of the gene target, before hypothesis testing and ecological meaning can be inferred from amplicon data sets, the sequences first have to be grouped into taxonomically meaningful ‘units’ to allow downstream analysis. There are two approaches to group amplicon data for downstream analysis - Operational Taxonomic Units (OTU) and Amplicon Sequence Variants (ASV). OTUs group sequences into a consensus sequence (the OTU) at a defined sequence similarity threshold. For the *16S rRNA* gene, a threshold of 97% sequence identity is generally used to define OTUs at species level, although this value has been challenged (18). As functional genes may have been subject to significant horizontal gene transfer, and be present in poly or monophyletic groups, the relationship between percentage identity and taxonomic delimitation is not clear and often unknown. As a result, in the literature different similarity thresholds have been used to construct OTUs for the same functional gene target (Table 1), with an OTU at 97% widely used, often without clear rationale. This variation in OTU selection criteria makes it difficult to select a meaningful value and creates limitations when comparing among studies. This is important as uncertainties in selecting the appropriate taxonomic cut-off could lead to different interpretations of findings underpinning our understanding of larger scale ecological processes and mechanisms structuring microbial communities and their activities (19). For the *16S rRNA* gene, the choice of OTU similarity threshold used can significantly influence microbial diversity patterns (20, 21) and we hypothesize this is also true for functional genes. In fact, some authors suggested that OTU similarity thresholds should be adjusted depending on the clustering algorithm and data complexity when phylogenetically divergent groups are present within the same community. This is because a single threshold for species delimitation is often not relevant due to variable evolutionary rates of the *16S rRNA* gene across lineages (22, 23). Indeed, the use of strict thresholds has been shown to result in phylogenetically inconsistent (para- or poly-phyletic) OTUs (24, 25).

**Table 1.** OTU similarity thresholds used in the literature to construct OTUs from nitrogen-cycle genes.

|  | Primers | Similarity thresholds |  |  |  |  |  |
| --- | --- | --- | --- | --- | --- | --- | --- |
|  |  | 97% | 95% | 90% | 85% | 83% | 82% |
| **AOB *amoA***<br>**Bacterial**<br>***Ammonia monooxygenase*** | BacamoA-1F/<br>BacamoA-2R | (7, 8, 39–42) (43) | (44–46) | (47) | (48, 49) |  |  |
| **AOA *amoA***<br>**Archaeal**<br>***Ammonia monooxygenase*** | Arch-amoWAF/<br>Arch-amoWAR | (7, 8, 40, 41, 50, 51) (43) | (44, 46, 48) | (52) | (47–49) |  |  |
| ***nirK***<br>***Nitrate reductase*** | nirK FlaCu/<br>nirK R3Cu | (53–55) | (46) | (56) |  | (57) |  |
| ***nirS***<br>***Nitrate reductase*** | nirS1F/<br>nirS 3R | (54, 55) | (58) | (56) |  |  | (57) |
| ***nrfA***<br>***nitrate reductase to ammonia*** | nrfAF2aw/<br>nrfAR1 | (59) |  |  |  |  |  |
| ***nxrB***<br>***nitrite oxidase*** | nxrB169f/<br>nxrB638r | (60) |  |  |  |  |  |

After selection of the sequence similarity threshold, further choices need to be made. OTUs clustering can be done using a closed-reference approach (22, 26) or by *de Novo* assembly (22, 26). With the latter method, reads are clustered into OTUs without comparison to a pre-existing database. This helps with the discovery of novel OTUs that are dissimilar to known sequences. However, the absence of comparison to known references makes *de novo* OTUs data-set-dependent, meaning that *de novo* OTUs from two different studies cannot be directly compared. On the other hand, the former approach makes it easier to compare OTUs between studies but limits the possibilities of discovering new sequence (26).

With the Amplicon Sequence Variant (ASV) approach (26), an error model is generated for the sequencing run, and reads are clustered in order to map this error model. This approach is appealing as it no longer groups amplicons based on a consensus sequence but instead resolves sequences with as little as a single nucleotide variation. Consequently, the ASV method does not require reference databases, is able to detect new sequences and ASVs from different datasets can be directly compared (26). The ASV method has been compared to the OTU approach using phylogenetic marker (*16S rRNA*, *18S rRNA* and fungal ITS) and overall results indicate better accuracy (27–29) and sensitivity (30) of the former when tested against mock communities. The impact this has on large scale ecological patterns still needs to be fully understood as some research suggests that the biological conclusions drawn using either method, based on phylogenetic markers, are largely consistent (28, 31) while other suggest that it can affect the interpretations of differentially expressed taxa between treatments (32). Nonetheless, as the use of ASVs doesn’t rely on a user-defined threshold that may not hold biological meaning, this approach should increase phylogenetic resolution of functional genes and, importantly, facilitate comparison among studies, as they should be able to segregate sequences on as little as one nucleotide variant. Consequently, ASVs are a promising approach for functional gene amplicon studies (13). However how the use of ASVs versus OTUs impacts ecological interpretation based on functional genes is unclear.

After selecting the appropriate amplicon reconstruction method (ASV or OTU) the next immediate challenge is assigning taxonomy by matching against a reference database. For *16Sr RNA*, several curated databases are available (*e.g.* Silva, GreenGene, Midas) and are routinely used. For most functional genes, no such databases are available and the reference sequences used to assign taxonomy often vary among studies. The most popular approaches rely on pattern recognition of overlapping “words” of length k (generally k=8) called k-mer. The frequency of matching k-mers between query and reference sequences is used as a measure of sequence similarity: higher frequency of shared k-mers indicates higher similarity between query and reference. This approach is fast, objective, and not limited by the uncertainties associated with methods based on evolutionary models and alignments (33, 34). This approach is usually implemented as a classifier, such as the commonly used Naïve Bayesian Classifier (NBC). One main limitation of this approach is that it uses the assumption that the actual position of the k-mers in the sequence is not important whereas, in reality, two sequences with the same k-mer but in different orders are different. Furthermore, the optimal choice of length for the k-mer might vary depending on the target gene or the region within the same gene (35). Another approach is the Bayesian Lowest Common Ancestor (BLCA) (35) where the query sequence is BLASTed against a reference database(s) and significant “hits” recorded. The taxonomy of the query sequence is assigned as the lowest common ancestor between these hits. For example, if a query sequence has significant matches to two *Nitrosomonas europaea* references sequences, the query sequence will be assigned to the species *Nitrosomonas europaea* as it is the lowest common ancestor between the two hits. However, if a query sequence has two different hits: *Nitrosomonas europaea* and *Nitrosomonas oligotropha*, the query sequence will be assigned to the genus *Nitrosomonas*. By considering hit results from multiple databases, the BLCA approach is able to provide probabilistic-based confidence values at each taxonomic levels of this assignment. Previously, the BLCA method has been shown to provide better species-level resolution than the NBC for *16S rRNA* sequences (35, 36). How they compare for functional genes has yet to be determined.

The aim of this study is to compare the effect of amplicon reconstruction approaches: OTUs (at a range of sequence similarity values) vs. ASVs and the taxonomic assignment method, NBC vs. BLCA, on a suite of functional genes. We do this to determine if diversity measures and subsequent ecological interpretation are affected by these choices. We hypothesize that both alpha and beta diversity measures will differ depending on the amplicon processing methods used. To do this, we examine the nitrogen cycle in marine sediments of Marennes-Oléron bay, French Atlantic coast. The middle part of the bay, the Brouage mudflat, is characterized by the presence of flow-parallel sediment structures consisting of crests (ridges) and troughs (runnels). These side-by-side physical structures have been shown to significantly affect nitrification rates (higher in runnels) (37, 38). We ask if the physical structure of the ridges and runnels results in differences in the diversity of the nitrogen cycling community present. Different pathways of the nitrogen cycle are targeted via genes encoding key enzymes. Specifically, nitrification, the oxidation of ammonia to nitrate via nitrite, is targeted via the subunit A of the ammonia monooxygenase *amoA*, and the beta subunit of nitrite oxidoreductase *nrxB*; denitrification, the sequential reduction of nitrate to di-nitrogen gas, via the nitrite reductase genes *nirS* and *nirK* and dissimilatory nitrate reduction to ammonia (DNAR) via cytochrome C nitrite reductase *nrfA*.

## Material and Methods

- Similarity thresholds from the literature used to cluster OTUs for nitrogen-cycle-genes (See Table 1)
- Sample collection, physiochemical measurements, PCR and Illumina sequencing

Triplicate surface mud cores (0 to 2 cm) were sampled July 2016 from the Montportail-Brouage mudflat, France, from three different ridges and runnels within 27.41 m^2^ (location 1 (L1): 45 54 31.50 N; 001 05 14.60 O/ Location 2 (L2): 45 54 31.70 N; 001 0 514.20 O/ Location 3 (L3): 45 54 31.50 N; 001 05 14.20 O) (61, 62) at the VASIREMI station (Figure 1). Sediment was homogenised and collected in sterile 5ml syringes, flash frozen and stored at -80°C until subsequent use. Bio-physio-chemical parameters were measured as described in (62), detailed procedures are provided in Supplementary Methods. DNA extraction was carried out using a modification of the protocol developed in (63), PCR of the *amo*A (bacteria and archaea), *nrxA*, *nirS/K* and *nrfA* was carried out as detailed in Table 2. Illumina Amplicon sequencing library preparation was carried as described in (10) using the Nextera XT Index Kit (Illumina, UK). Products were pooled at equimolar concentration and submitted to Earlham Institute (Norwich Research Park, Norwich, UK) for Illumina MiSeq sequencing (300PE, 22 millions reads/lane). Detailed protocols are provided in Supplementary Methods.

**Figure 1.**
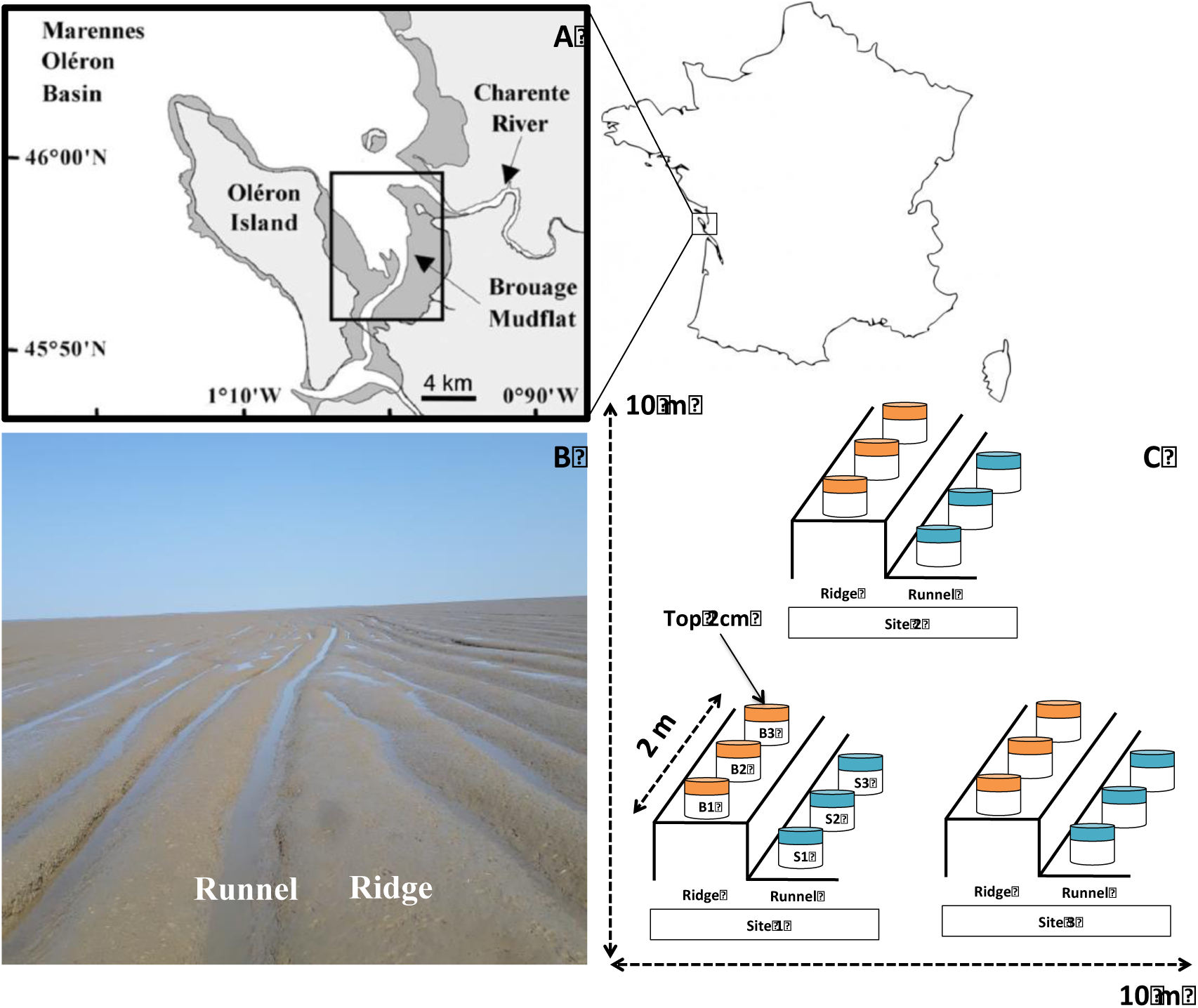
Sediment sampling of ridges and runnels on the Brouage mudflat. A) ap showing the location of the Brouage mudflat on the Marennes-Oléron Bay, French Atlantic coast. B) Parallel ridge/runnels sedimentary structures that characterise the intertidal mudflat. C) A schematic of the sampling plan; three ridge-runnels within ∼25m^2^ were sampled. For each ridge-runnel structure, replicate (n=3) sediment cores (2 cm depth) were taken form the ridges and runnels along a ≈ 2m transect.

**Table 2.**
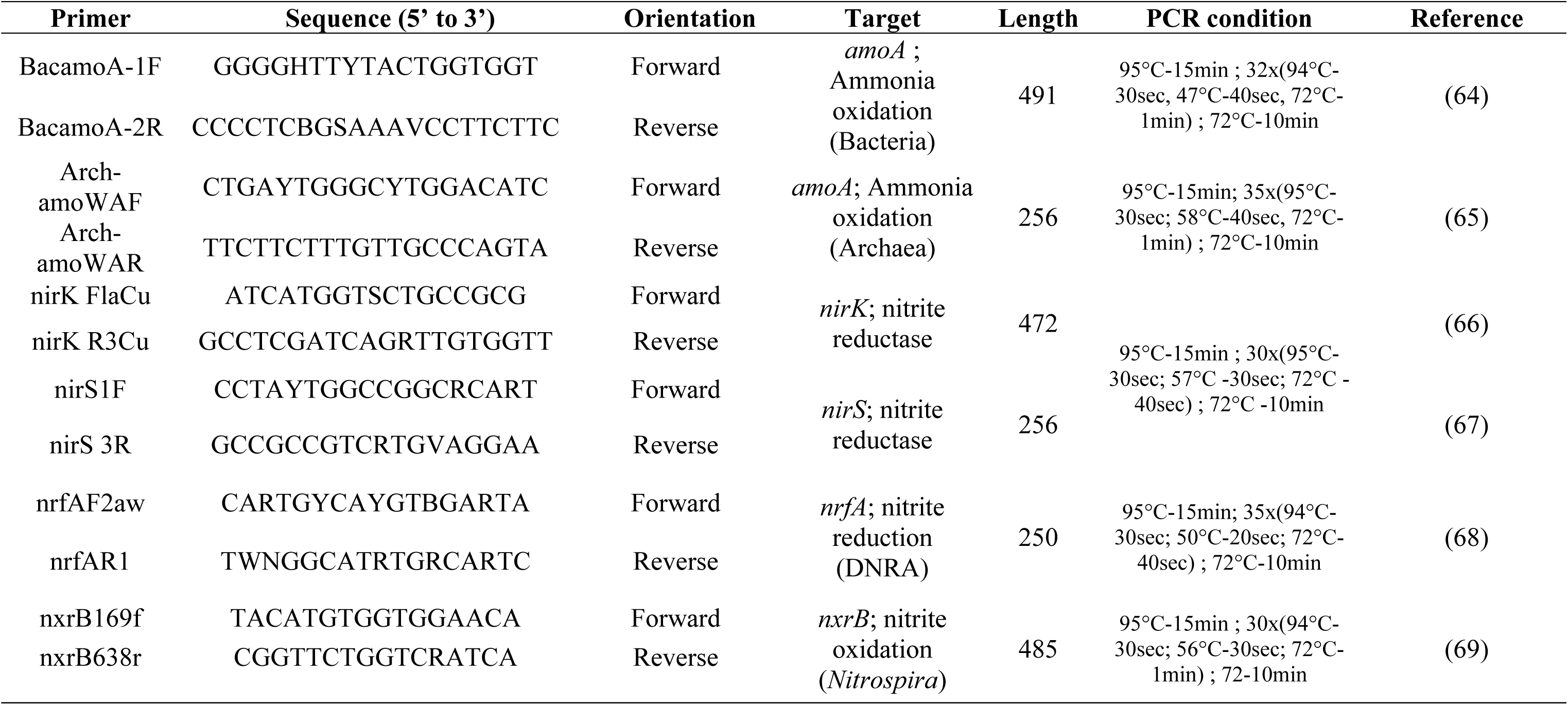
List primers and corresponding PCR conditions used in this study.

### OTU and ASV construction

To construct OTUs, paired-end reads were trimmed and filtered with Sickle v1.2 (70) using a sliding window and trimming regions where the average base quality was below 20. A 10bp threshold was used to discard reads below this length. BayesHammer (70), Spades v2.5.0 assembler was used to error correct paired-end reads followed by pandaseq v(2.4) with a minimum overlap of 20bp to assemble the forward and reverse reads into a single sequence. The choice of software was as a result of author’s recent work (71, 72) where it was shown that the above strategy of read trimming followed by error correction and overlapping reads reduces the substitution rates significantly. After having obtained the consensus sequences from each sample, the VSEARCH (v2.3.4) pipeline (all these steps are documented in https://github.com/torognes/vsearch/wiki/VSEARCH-pipeline) was used for OTU construction. Reads were pooled from different samples and barcodes added to keep an account of the samples the reads originated from. Reads were then de-replicated and sorted by decreasing abundance and singletons discarded. In the next step, the reads were clustered based on different similarity threshold (97%, 95%, 90% and 85% for AOA and AOB *amoA*; 97%, 95%, 90% and 83% or 82% for *nirK* (83%) and *nirS* (82%); 97% for *nxrB* and *nrfA*), followed by removing clusters that have chimeric models built from more abundant reads (`--uchime_denovo` option in vsearch). A few chimeras may be missed, especially if they have parents that are absent from the reads or are present with very low abundance. Therefore, in the next step, we use a reference-based chimera-filtering step (`--uchime_ref` option in vsearch) using the above created reference databases. The original barcoded reads were matched against clean OTUs with the different similarity thresholds to generate OTU tables.

To construct Amplicon Sequence Variants (ASVs) the DADA2 pipeline (73) was used on Qiime2, the full pipeline can be found at https://github.com/umerijaz/tutorials/blob/master/qiime2_tutorial.md. Briefly, reads were demultiplexed using `qiime demux emp-paired()` and denoised/quality trimmed using `qiime dada2 denoise-paired()`. For AOA *amoA*, *nirS* and *nrfA* HTS data, forward reads were trimmed at 240bp and reverse reads at 200bp. ASVs were then constructed by merging the forward and reverse ASVs together and dereplicated to generate abundance files (ASVs counts in each samples), with both process internal to DADA2 algorithm. To find the phylogenetic distances between ASVs, they were aligned using Maftt (74) and the phylogenetic tree was constructed using FastTree (75). For the Bacterial *amoA* and *nxrB* and *nirK*, datasets forward and reverse reads did not merge properly when using the previous method. Therefore, demultiplexed reads were processed in R using R’s DADA2 package (73). First, quality trimming was done using `filterAndTrim`(). Forward reads were trimmed at 275bp and 230bp, 213bp and 215bp for the reverse reads of *amoA* and *nxrB* and *nirK* respectively, allowing for a maximum of 2 errors in the forward and reverse reads to merge. Error models were generated using `learnErrors`(). Reads were dereplicated using `derepFastq`() and ASVs were inferred using `dada`(). Forward and reverse reads were merged using `mergePairs()`allowing for a minimum overlap of 10bp. A sequence table was generated using `makeSequenceTable()`and chimeras were removed using `removeBimeraDenovo()`. A count table was then generated and distances between the representative ASVs were inferred by aligning the sequences using Maftt (74) and constructing a phylogenetic tree using FastTree (75). All R scripts used for analyses are available at http://userweb.eng.gla.ac.uk/umer.ijaz/bioinformatics/ecological.html and as part of R’s microbiomeSeq package http://www.github.com/umerijaz/microbiomeSeq or in the supplementary material and methods.

### Taxonomic assignation: NBC and BLCA

For each nitrogen cycle gene, reference sequences (nucleotide) were downloaded from Fungene (http://fungene.cme.msu.edu/); for AOA and AOB *amoA*, a second database was constructed by downloading the sequences corresponding to the different clusters defined in Zhang *et al*, (8). Subsequently, R’s rentrez package (76) was used to get taxonomic information at different levels generating a taxonomy file. The FASTA file and the corresponding taxonomy file were formatted to work with Qiime (77).

To assign taxonomy to the representative ASVs, two different approaches were used: Representative ASVs were classified using a Naïve Bayesian Classifier K-mer classifier (NBC) (`qiime feature-classifier classify-sklearn`) or the Bayesian Lowest Common Ancestor (BLCA) (35) against the reference databases. A detailed protocol for BLCA can be found at https://github.com/qunfengdong/BLCA.

Count tables -generated in the previous step- and taxonomy tables were combined together to generate biom files using Qiime (77) (https://qiime2.org) (`biom add-metadata`) and the phyloseq package was used to load these biom files in R.

### Amplicons reads quality check

After OTUs and ASVs construction, a quality check step was undertaken to ensure the reliability of the data using R seqinr package (78). First, OTUs/ASVs nucleotide sequences were filtered based on their length, for AOA *amoA*, AOB *amoA*, *nxrB*, *nirS*, *nirK* and *nrfA*, nucleotide sequences that were <256bp, <491bp, <480bp, <250bp, <470bp, and <250bp respectively were deleted from the dataset. These “error-prone” sequences were checked using BLASTn on NCBI to ensure they were not genuine sequences. OTUs/ASVs were then translated to protein using MEGA7 (79) and BLASTed (BLASTp). OTUs/ASVs that did not translate to the correct protein or included stop codons were deleted from the dataset.

OTUs and ASVs sequences were then clustered based on amino acid sequence similarity at 100% similarity threshold: OTUs/ASVs that translated to the same protein sequence were considered the same and their abundances were added together in the abundance files (Supplementary Method). The final abundance file therefore contained only correct sequences that translated to unique proteins (Figure 2). Finally, a phylogenetic tree was constructed using these correct and unique OTUs/ASVs: DNA sequences were aligned using Mafft and the phylogenetic trees were obtained using FastTree on these alignments. The tree was visualised in R using the ggtree package (80). Downstream statistical analyses were carried out in R (see Supplementary Method for details).

**Figure 2.**
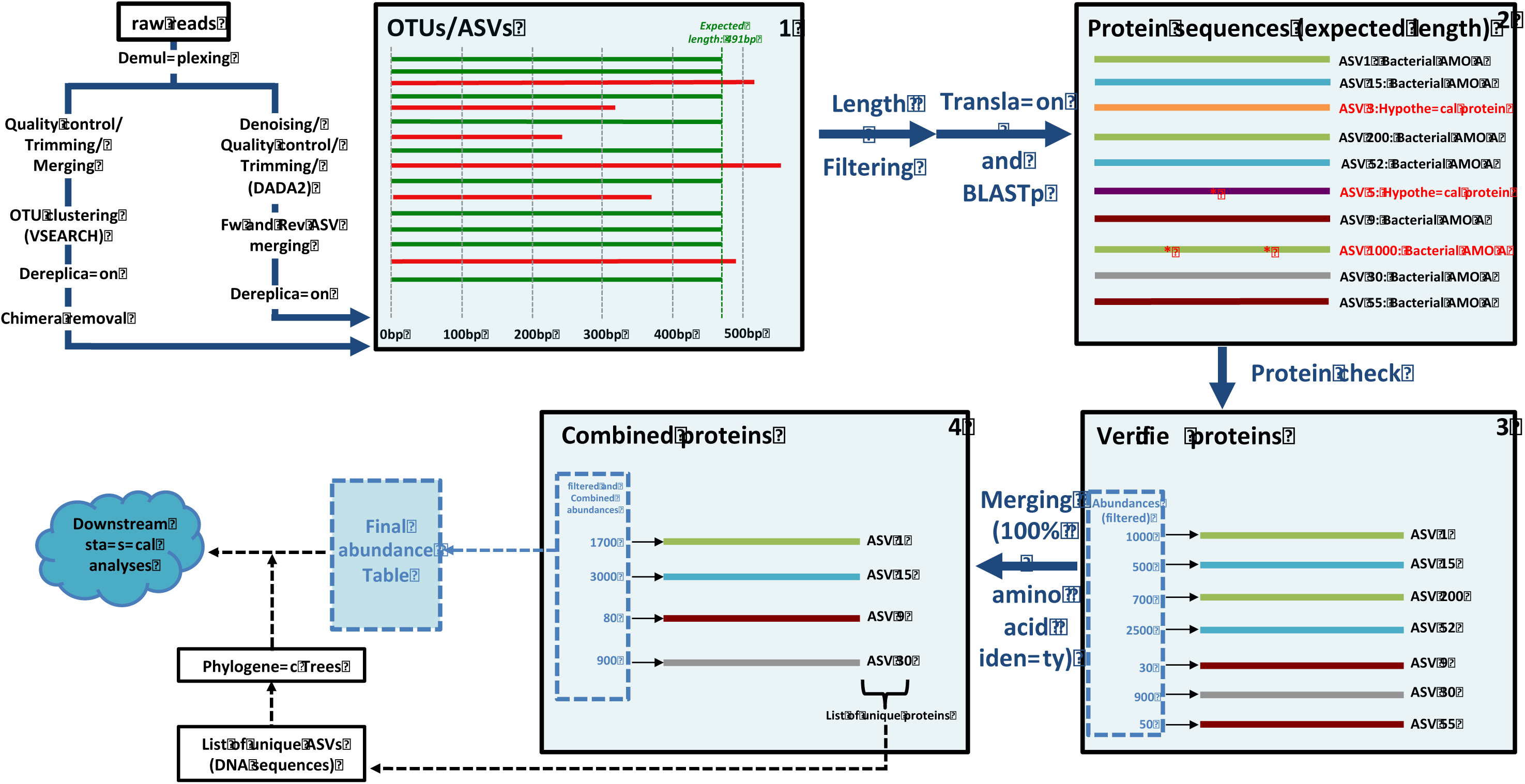
OTU/ASV quality check workflow (example of the Bacterial *amoA*). Red reads in step 1 are reads of the wrong length. Colour of the proteins in step 2 and 3 represent their amino acid sequences (two sequences with the same colour are identical); red asterisk indicates stop codons. Between steps 3 and 4, the abundances of identical proteins have been added together.

## Results

### Effect of OTU threshold and ASV selection on α diversity and sequence coverage

#### 1) *α* diversity indexes

Richness, Simpson and Shannon indexes were calculated based on the rarefied abundances tables in ridges and runnels obtained by using different percentage identity for OTU construction (Table 1) or using ASVs. In general, an increase in the percentage identity used to generate OTUs resulted in an increase in the value of alpha diversity indexes. A similar increase was generally observed when using ASVs instead of OTUs. The increase in alpha diversity indexes between OTUs and ASVs was particularly strong for *nirS* and *nrfA*. In contrary, for AOA *amoA* and *nxrB*, there was a slight decrease in Shannon and Simpson indexes from OTU-97% to ASVs. This was also observed for *nirK* but only in ridges. Interestingly, the use of OTUs (97%) and ASVs often lead to different conclusions as to which of the two sedimentary structures was the most diverse. This was especially true for *nirK* where the use of OTUs always resulted in higher values for all three indexes in ridges compared to runnels and inversely when using ASVs (Figure 3).

**Figure 3.**
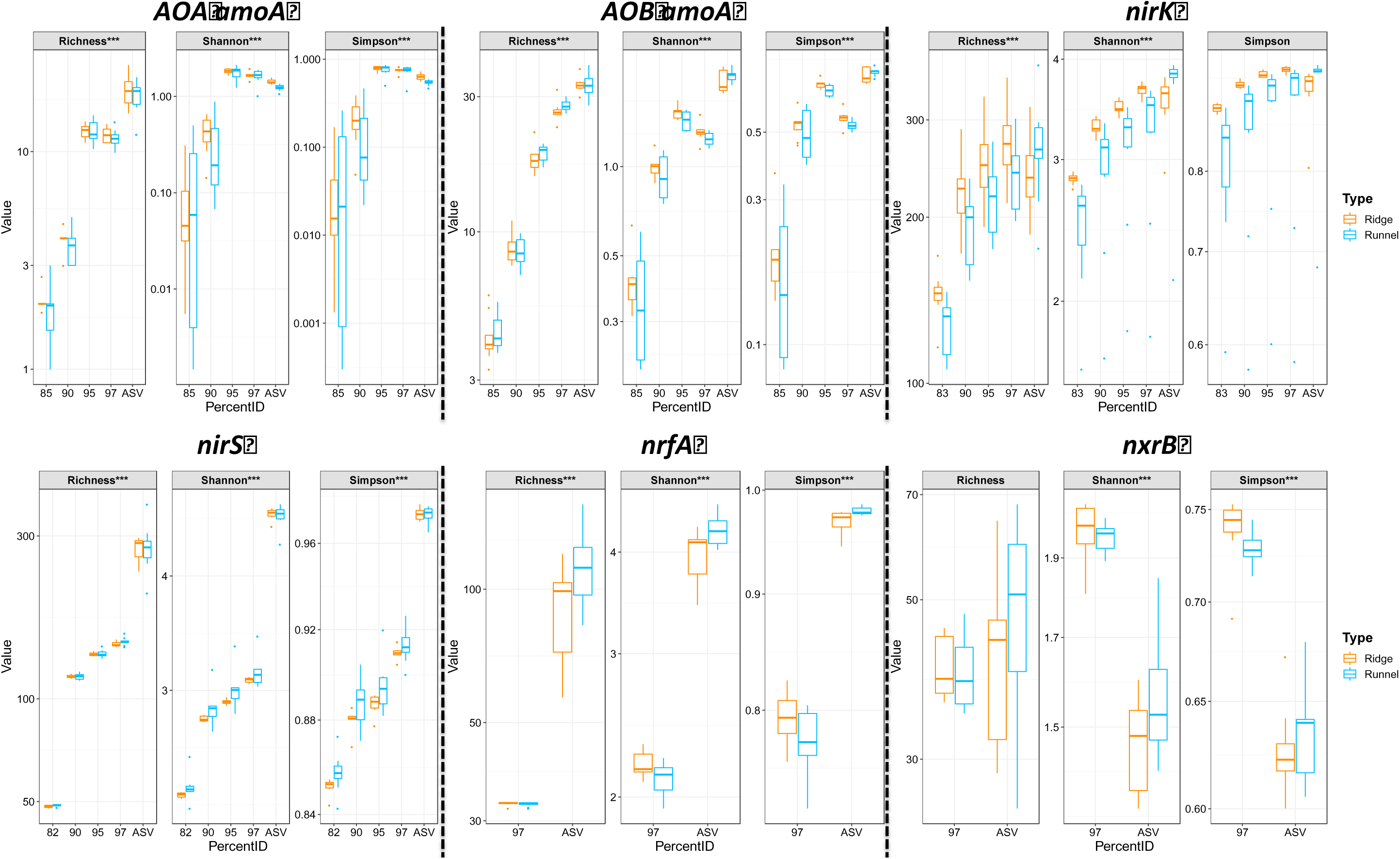
The effect of different OTU sequence similarity thresholds verses ASVs on *α* diversity results. Result of the ANOVA test for the effect of the clustering method on richness, Simpson and Shannon indexes is reported on top of each plot for each gene. *: *p-*value<0.05; **: 0.01>*p-*value>0.001; ***: *p-*value<0.001.

#### 2) Rarefaction curves

To determine if the sequencing effort had been sufficient to capture the full OTUs/ASVs diversity, rarefaction curves were drawn for all genes using the OTU and ASV abundance tables. When sequencing reads were clustered together using the OTU approach, the rarefaction curves generally reached a quasi-plateau phase, indicating that the observed richness was close to its maximum theoretical value. For *nirK* and *nxrB*, this plateau phase was not reached, indicating that more OTUs would have been recovered with greater sequencing depth. Rarefaction curves obtained from ASV abundance tables were generally further from reaching the plateau compared to curves obtained using the OTU approach. This was particularly true for *nirS*, *nirK* and the bacterial *amoA* genes (Figure 4).

**Figure 4.**
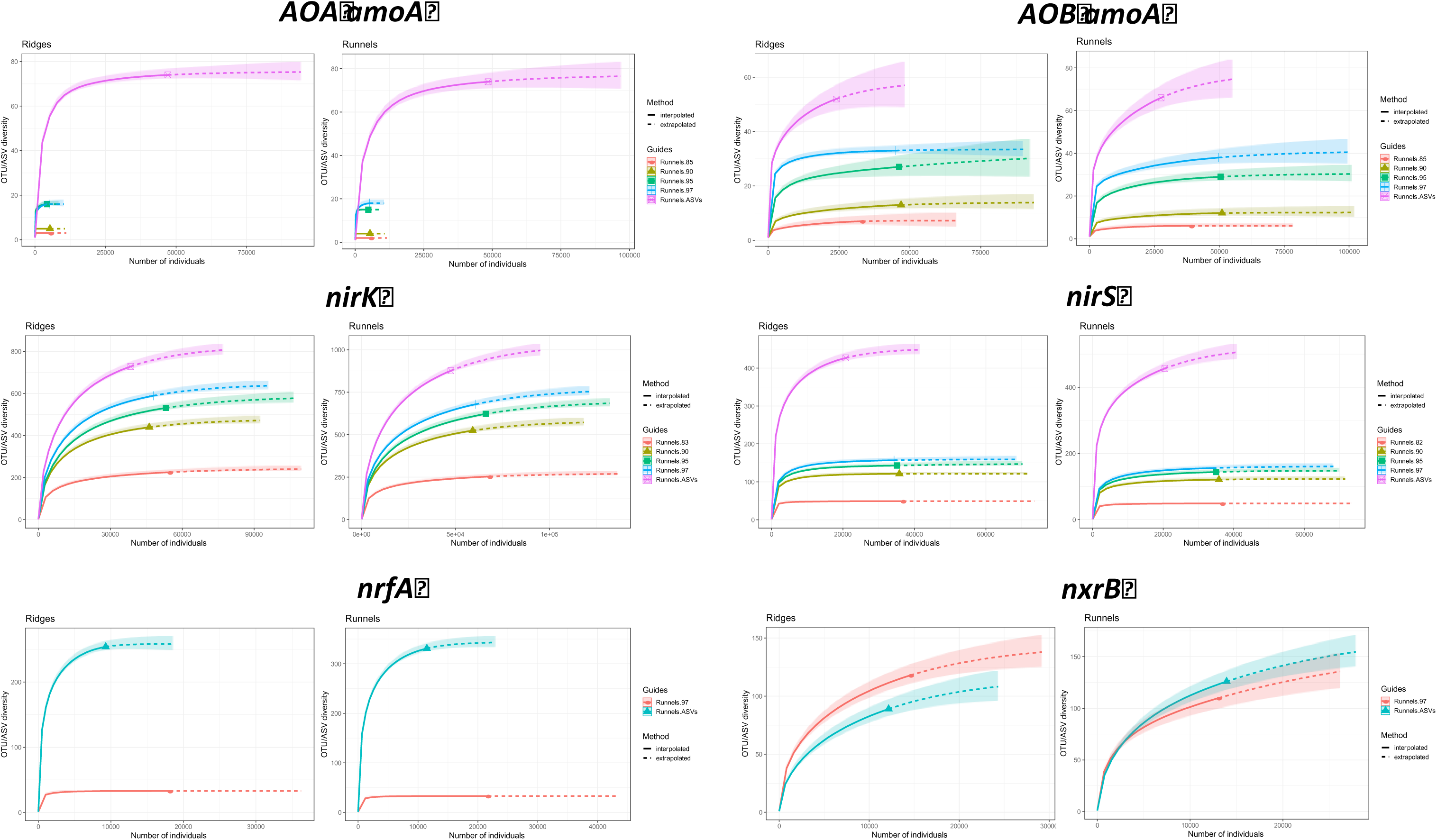
Rarefaction curves in ridges and runnels using different OTUs sequence similarity thresholds and ASVs.

### Effect of OTU thresholds and ASVs on β diversity

#### 1) Dissimilatory distances between samples

To determine the effect of the amplicon data analysis method on the dissimilarity distance between samples, Mantel correlations were calculated between distances matrices obtained using Bray-Curtis (BC), Unifrac (U) and Weighted Unifrac (WU) metrics. A strong effect of the approach used is seen as reflected by Mantel correlations different from 1 (Figure 5). For *nirK*, low correlation between distance matrices obtained with ASV vs. OTU were observed, while the correlations between distances matrices obtained with OTUs at different similarity threshold were generally high (>0.5; *p-*value<0.01). For *nirS*, a similar trend was observed especially when considering Unifrac distances. For AOA and AOB *amoA*, again, the correlations between distance matrices obtained with Bray-Curtis (and Weighted Unifrac for AOB) were always lower when considering ASV vs. OTU than for OTUs at different similarity threshold. Correlations values for the Unifrac distance matrices did not follow any obvious pattern for these two genes.

**Figure 5.**
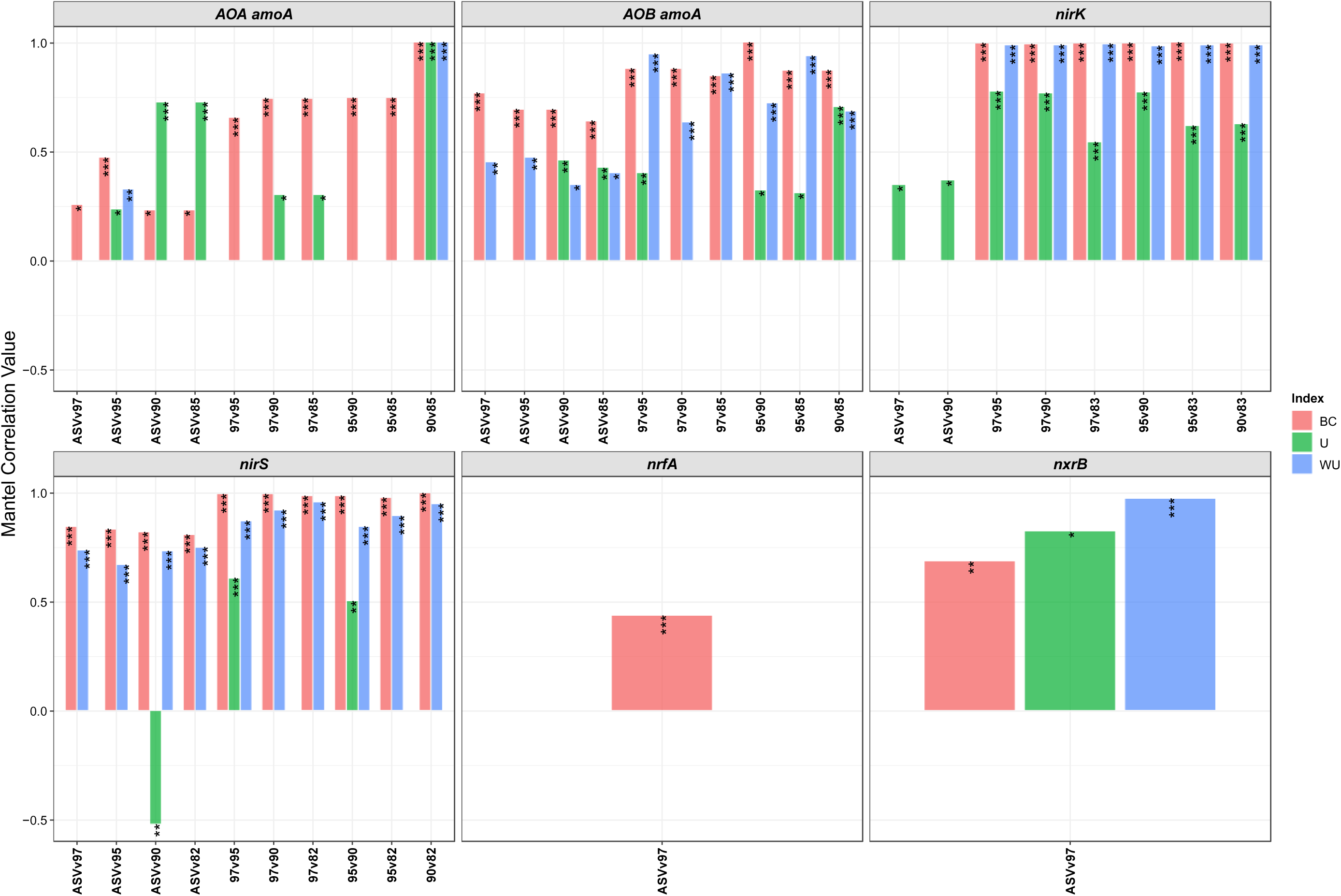
Effect of different OTU sequence similarity threshold or ASV amplicon reconstruction approach on phylogenetic distances. Distances matrices were calculated using Bray-Curtis (BC), Unifrac (U) and Weighted Unifrac (WU) metrics at different OTU similarity threshold and ASV for each gene. Pairwise correlations between matrices, as indicated on the x-axis, were calculated using Mantel test (only significant values are represented). *p-*value of the tests are reported on top of the barplot: *: *p-*value<0.05; **: 0.01>*p-*value>0.001; ***: *p-* value<0.001.

#### 2) Differences in communities between ridges and runnels

To determine whether the method used to process sequencing reads could significantly affect *β* diversity results, Bray-Curtis, Unifrac and WUnifrac distances between communities in the two sedimentary structures (ridges and runnels) were calculated and their significance tested using PERMANOVA. The effect of the clustering method was different depending on the gene of interest (Figure 6). For AOA *amoA*, significant differences in WUnifrac distances between ridges and runnels are seen when reads were clustered using the ASV but not when using OTUs, regardless of the percentage identity threshold used. Similarly, significant differences were observed for *nrfA* communities, for all three beta-diversity indexes when using ASVs but not when using OTUs. Interestingly, for AOB *amoA*, a significant effect of the sedimentary structure was observed on WUnifrac distances when using OTUs at 97% similarity threshold but not for other OTU values or for ASVs. For the other genes (*nirS*, *nirK* and *nxrB*), the different clustering methods were more consistent, with no significant effect of the data analysis pipeline on *β* diversity (Figure 6). To summarise, the clustering method used for amplicons had a significant effect on the beta diversity for some genes while other were less affected.

**Figure 6.**
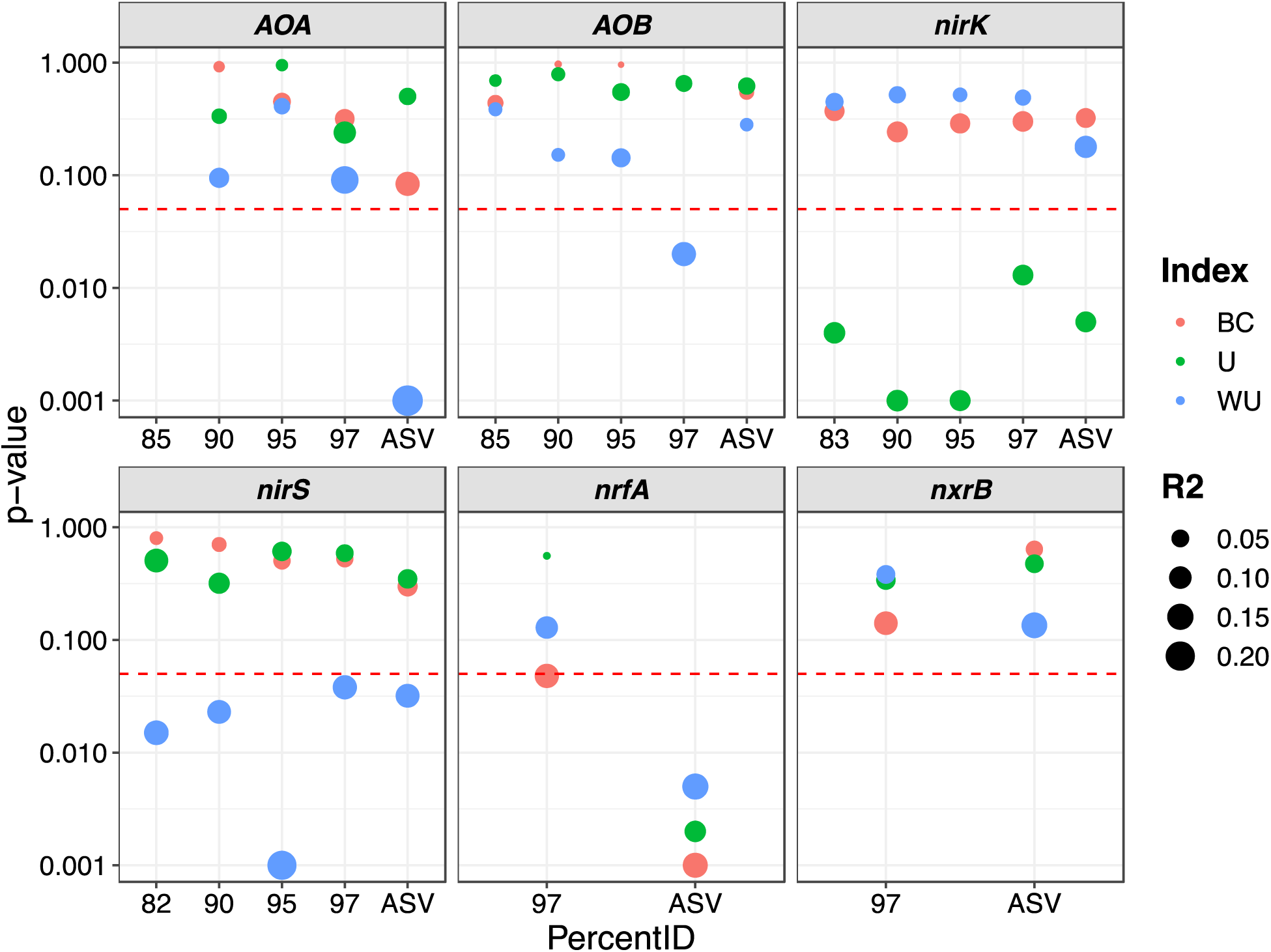
Effect of amplicon reconstruction method on *β* diversity results. The reported *p-*value shows the effect of the ridge/runnel structure on the community composition determined by the sequencing of six different nitrogen cycle genes using three different dissimilarity distance, indicated by the colour of the symbols as explained in the legend: BC=Bray-Curtis, U=Unifrac, WU=Weighted Unifrac. The horizontal dashed red line represents the p-value threshold for significance (0.05). The size of points represents the R squared, *i.e.* the percentage of variance in BC, U and WU explained by the ridges/runnels structures.

To better understand the effect of OTUs vs. ASVs on the phylogeny of the representative sequences, phylogenetic trees were drawn using representative OTUs/ASVs along with full-length sequences from isolated microorganisms downloaded from NCBI for genes where the HTS data analysis method had a significant effect (AOB *amoA* AOA *amoA* and *nrfA*). For clarity, a second set of trees was generated where all nodes leading only to OTUs/ASVs were collapsed (Supplementary Figure 1). For AOB *amoA*, the OTU and ASV collapsed trees shared a very similar internal structure. For AOA *amoA* and *nrfA*, the internal structures of the collapsed trees were more dissimilar.

### Effect of OTU thresholds and ASVs on canonical analyses

Canonical analyses, *e.g.* canonical correspondence analysis (CCA; http://www.github.com/umerijaz/microbiomeSeq), are useful approaches to analyse, detect and visualize interactions between microbial communities and environmental parameters. CCA measures the association between an explanatory table (the physiochemical parameters) and a response table (the abundance table). Previously, it was shown that, in some cases, the amplicon reconstruction method affects the composition of the abundance tables as revealed by Mantel correlations between Bray-Curtis distances matrices obtained from ASVs or OTUs at different similarity thresholds lower than 1 (Figure 5). These changes in the abundance tables can be expected to also change the results for CCA. To test this, CCA was done using the abundances tables obtained from the different amplicon reconstruction methods (range of OTU sequence similarity thresholds and ASVs) using the same physiochemical data table. As seem in Table 3, the choice of amplicon processing methods strongly impacted results. For ammonia oxidisers, when using ASVs none of the measured physiochemical parameters measured affected AOB but nitrate was a significant driver of AOA. Interestingly, for both ammonia oxidizer communities, the highest number of environmental drivers was found when OTUs were clustered at 95% similarity threshold. TOC was a driver of both AOA and AOB communities but only at 97% and 95% whereas the sediment grain size (SGS) was a driver only at 90% and 85%.

**Table 3.** Canonical Correspondence Analysis. ChlA: chlorophyll A; SGS: sediment grain size (%<63µm); TOC: total organic carbon; DOC: dissolved organic carbon; TDN total dissolved nitrogen. Empty cells indicate that the physiochemical parameter is not a driver of the community.. :0.1>*p-*value >0.05; *: *p-*value<0.05; **: 0.01>*p-*value>0.001; ***: *p-*value<0.001.

|  | NH<sub>3</sub> | NO<sub>2</sub><sup>-</sup> | NO<sub>3</sub><sup>-</sup> | ChlA | pH | SGS | TOC | DOC | TDN |
| --- | --- | --- | --- | --- | --- | --- | --- | --- | --- |
| *AOB ASV* |  |  |  |  | . |  |  |  |  |
| *AOB 97* |  |  |  |  |  | . | ** |  | . |
| *AOB 95* | . |  | * |  | ** |  | * |  | . |
| *AOB 90* |  |  |  |  | . | * | . |  |  |
| *AOB 85* |  |  |  |  |  | * | * |  |  |
| *AOA ASV* |  |  | * |  |  |  |  |  |  |
| *AOA 97* |  | * | . |  | . |  | ** |  |  |
| *AOA 95* | ** | * |  | * | . |  | * |  |  |
| *AOA 90* |  |  |  |  |  | ** |  |  |  |
| *AOA 85* |  |  |  |  |  | * |  |  |  |
| *nirK ASV* | ** |  |  | * |  | ** | * |  |  |
| *nirK 97* |  |  |  |  |  | * |  |  |  |
| *nirK 95* | * |  |  | . |  | * |  |  |  |
| *nirK 90* |  |  |  |  |  |  |  |  |  |
| *nirK 83* |  |  |  |  |  |  |  |  |  |
| *nirS ASV* |  |  |  |  | . | * | . |  |  |
| *nirS 97* |  |  |  |  |  |  |  |  | * |
| *nirS 95* |  |  |  | . | * |  |  | . |  |
| *nirS 90* |  |  |  |  |  |  |  |  | . |
| *nirS 82* |  |  |  | . |  |  |  |  | * |
| *nxB ASV* | . |  | ** |  |  |  |  | . | * |
| *nxB 97* |  |  |  |  |  |  |  | * |  |
| *nrfa ASV* |  |  |  |  | ** |  |  |  |  |
| *nrfa 97* |  |  | * | ** | ** |  |  |  |  |

For the nitrite oxidising community, again, differences were seen depending on amplicon reconstruction approach. DOC was found to be a driver of the nitrite oxidizers communities but only with OTUs 97% and this effect became non-significant with ASVs. The opposite was true for nitrate and TDN, which were significant with ASVs only.

For denitrifiers, *nirK*, sediment grain size (SGS) was a strong driver (ASV, OTU 97% and 95%), along with ammonia (ASV and OUT 95%) and, to a lower extent chlorophyll A (ASV) and TOC (ASV). Generally, more physiochemical parameters were found to be significant drivers when using ASVs compared to OTUs. More significant drivers were generally found when using higher similarity thresholds for OTUs. For *nirS*, SGS was also a driver but only when using ASVs. TDN was a driver with OTU at 97% and 82% and pH with OTU at 95%. For dissimilatory nitrate reduction to ammonia, OTUs and ASVs for *nrfA* and pH were correlated whereas chlorophyll A and nitrate were only important when using OTU at 97%.

### NBC vs. BLCA for taxonomic classification

For the majority of genes studied here, neither the NBC nor the BLCA performed well, with a majority of ASVs unassigned at the species level when using the Fungene database. This is likely an indication of the lack of environmental sequences with species level-defined taxonomy in this database. The only exception was for *nrfA*, with NBC approach resulting in only 18.8% unassigned versus the BLCA in which 98.8% were unassigned. Both NBC and BLCA performed better when a custom database was used to assign taxonomy (AOA and AOB *amoA*). In this case, BLCA performed slightly better than NBC for AOA *amoA* and inversely for AOB *amoA* (Table 4). To determine what caused these differences for AOB *amoA*, a phylogenetic tree was constructed with AOB *amoA* ASVs and sequences from known AOB isolates downloaded from NBCI. The main differences in the taxonomic assignment for AOB *amoA* ASVs was for some sequences assigned as *Nitrosomonas aestuarii* with NBC that were unassigned using BLCA (Figure 7).

**Table 4.**
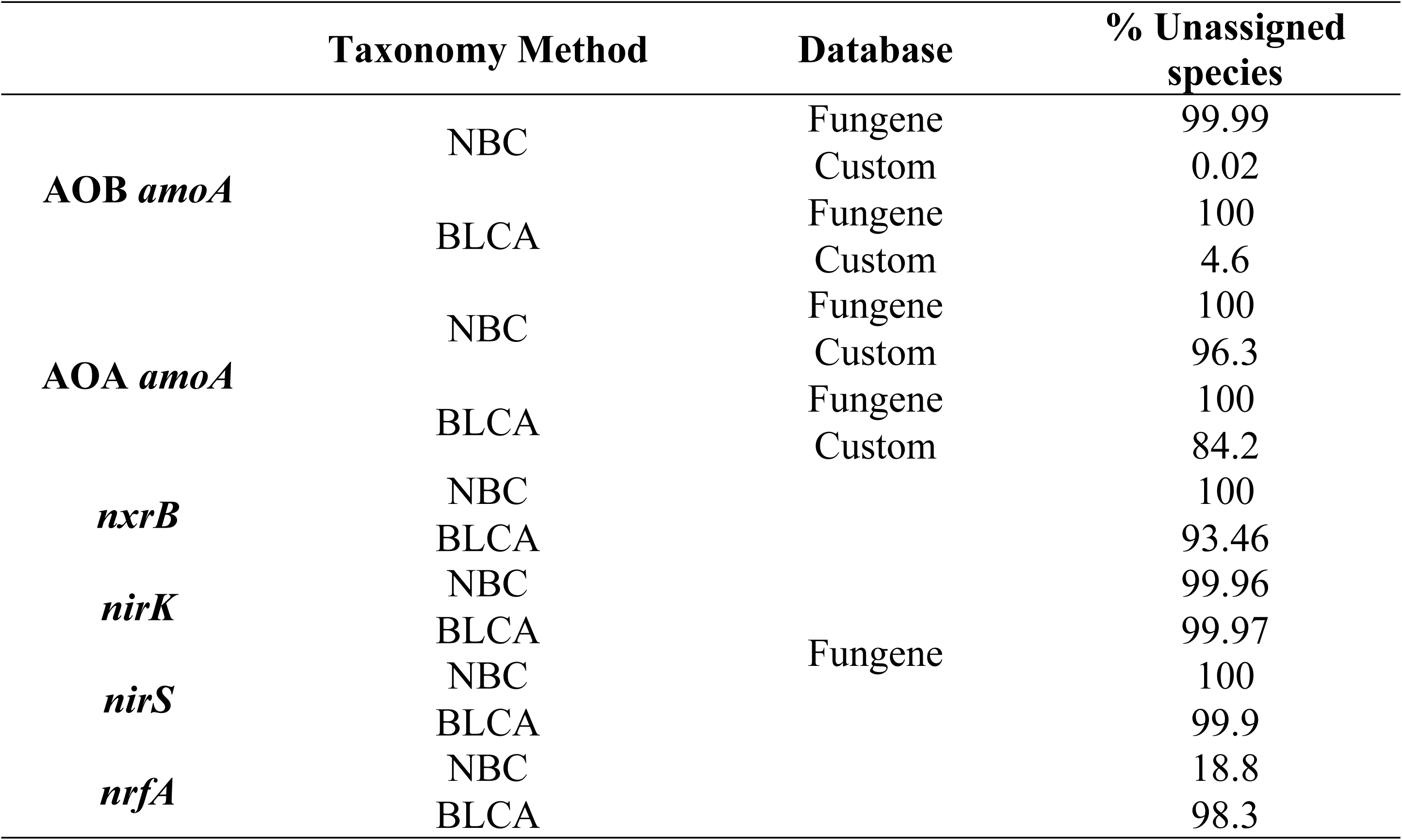
Performance of the NBC and BLCA methods for taxonomic assignment.

**Figure 7.**
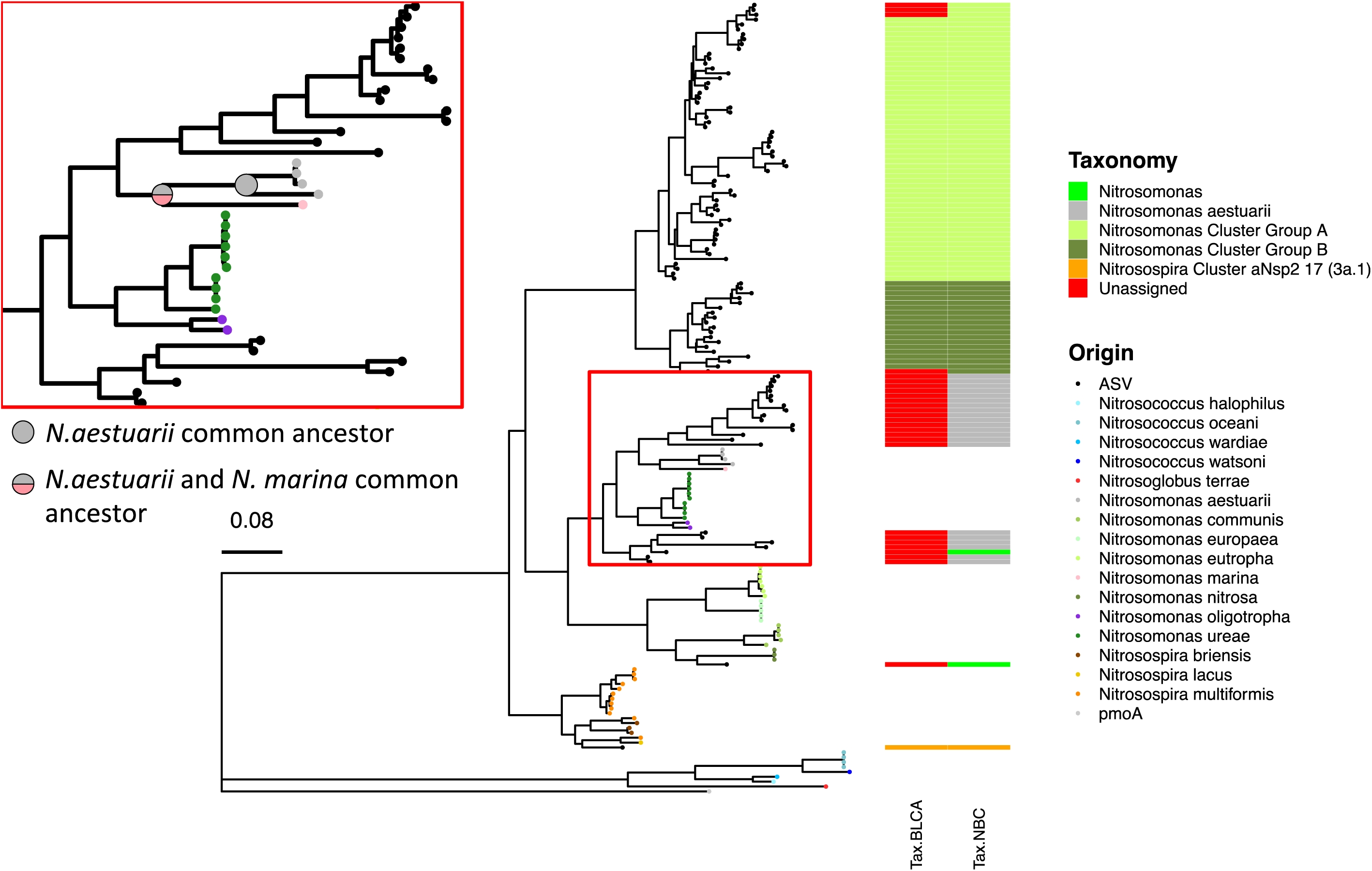
NBC vs. BLCA for the taxonomic assignation of AOB *amoA* ASVs. The taxonomy of representative sequences is indicated by the colour of the tips on the tree. Taxonomy assigned by NBC (TAX.NBC) and BLCA (TAX.BLCA) for each ASV is represented on the right of the tree. The main point of disagreement between NBC and BLCA is shown by a red rectangle (see zoom on the top left).

## Discussion

The aim of this study was to evaluate the effect of selecting different OTU similarity thresholds versus the ASVs approach for amplicon-sequence data processing. This was tested using a suite of functional genes targeting different pathways of the nitrogen cycle. We determined the effect these choices had on *α* and *β* diversity and on subsequent CCA analysis. We also examined the effect of the approach taken for taxonomic assignment of functional genes using the nitrogen cycle communities from ridges and runnels of the Montportail-Brouage mudflat as case study.

The choice of amplicon reconstruction method had a significant effect on biological observations (Figure 3, 5 and 6). As a result, the conclusions of this study based on OTUs (with different similarity threshold) or ASV leads to different ecological conclusions. For the AOA (WUnifrac distances) and *nrfA* communities (all distances), using the ASV denoising approach, we conclude a significant effect of the ridge/runnel sedimentary structures but no effect for AOB. On the other hand, based on OTU clustering at 97% similarity we conclude a significant effect of the ridge/runnel structures on AOB (WUnifrac). The results from this study show the strong effect the processing method has on the interpretation and biological understanding of sequencing data. This further illustrates the need for a standardised protocol for amplicon data processing to facilitate comparison between studies. To date, there is no consensus as to which threshold to use to construct OTUs for the same gene amplified with the same primers (Table 1) and we therefore argue that ASVs, which do not require user-defined similarity thresholds, offer a better chance to achieve such standardisation. We further showed that the choice of amplicon reconstruction method affects the outcome of multivariate analysis, which are routinely used to inform associations between biological assemblages and environmental parameters. For example, research on nitrification in the environment often seeks to determine the extent to which ammonia, pH, salinity, temperature etc. are significant drivers of the niche differentiation between AOA and AOB (7, 8, 88, 9, 81–87). This study clearly shows that the influence of environmental parameters on the ammonia oxidising communities is dependent on the amplicon reconstruction method used. A consensus standardised method needs to be adopted in molecular microbial ecology to allow for meta-reviews of current literature and the identification of ecological patterns that are not study-dependant.

To understand the effect of the choice of ASV versus OTU on phylogenetic resolution, phylogenetic trees were constructed from ASVs and OTUs (97%) for AOB *amoA*, AOA *amoA* and *nrfA* (genes for which the use of OTU vs. ASV had a significant effect). For AOB *amoA* it seemed that the change between OTUs and ASVs did not result in an overall different phylogenetic tree (Supplementary Figure 1). The ASVs tree is more detailed in terms of the number of tips (greater taxonomic resolution) than the OTU tree, which is clustering more of these sequences (tips) together, leading to a lower resolution of AOB diversity. However, for AOA *amoA* and *nrfA*, the trees obtained using either OTUs or ASVs were more dissimilar indicating that the OTU approach may cluster sequences that are different resulting in different phylogenies. This could be an effect of the shorter amplicon size for AOA *amoA* and *nrfA* compared to AOB *amoA*. Indeed, it has been shown before, for the *16S rRNA* gene, that the similarity threshold for short amplicons might differ from that of the full-length genes, resulting in inappropriate OTU clustering (89, 90). For AOB *amoA*, the amplicon covers more than half of the full-length gene, which might explain why its taxonomy is more consistent between OTU and ASV.

Other studies have reported that for *16S rRNA* amplicon sequencing, ecological patterns are robust to the choice of OTU vs. ASV(28, 31, 91). Similar results were observed in this study for the two nitrite reductase genes *nirS* and *nirK* where the use of OTUs at different thresholds or ASV led to the same biological conclusions: differences in composition between ridges and runnels when considering weighted Unifrac and Bray-Curtis distances for *nirS* and *nirK*, respectively (Figure 6). Interestingly, these two genes were the one with the highest richness (Figure 3). This is to be expected because these polyphyletic genes are widely spread throughout bacteria (92). In contrast, *amoA* encodes the ammonia monooxygenase and is restricted to nitrifiers. It could therefore be hypothesised that the differences between the *nir* genes (and *16S rRNA* as reported elsewhere) and the other functional genes is due to a resolution effect: when targeting a high diversity gene such as the *16S rRNA* or *nir* by amplicon sequencing, we obtain an overall picture of the microbial community. Therefore, the use of OTUs (low resolution) or ASV (high resolution) matters less because we still see the overall trends in the bacterial community. However, when investigating a phylogenetically tight group *e.g. amoA*, we are already zoomed in on a small part of that microbial community which explains why we need higher resolution *i.e.* ASVs to differentiate each members of the community.

This study also provides a comparison between NBC and BLCA for taxonomic assignment of functional gene sequences. We found that the BLCA method performed better than the NBC for AOA *amoA* when using a custom database and inversely for AOB *amoA* due to some sequences being assigned to *Nitrosomonas aestuarii* when using NBC and unassigned when using BLCA. When comparing these sequences to known representatives, it was observed that known AOB *amoA* sequences from *Nitrosomonas aestuarii* clustered together and the common ancestor for all *N. aestuarii*, did not include any of these ASVs. In fact, based on this tree, *N. aestuarii* and *N. marina* share a common ancestor that doesn’t include any ASV sequences. Based on this phylogeny, these ASVs could therefore not be members of the *N. aestuarii* group (Figure 7). We conclude that, for AOB *amoA*, the BLCA method, despite resulting in a lower number of sequences identified to species level, resulted in a classification that made more sense from a phylogenetic point of view. This result is coherent with previous research showing the superiority of BLCA vs. NBC for the taxonomic assignment of *16S rRNA* reads (36). However, the BLCA method performed much worse than NBC for the taxonomic assignation of *nrfA* sequences with the vast majority (98.8%) of unassigned sequences versus 18.8% for NBC. This likely reflects limitations in the database used rather than a problem with the BLCA method itself. Indeed, the majority of *nrfA* sequences in the Fungene database are full-length sequences originating from cultured microorganisms, which likely differ from sequences retrieved from the environment. To test this hypothesis, a phylogenetic tree was drawn using the top 50 *nrfA* ASVs found in this study and sequences from the reference database (covering 125 different genera). ASV sequences and representatives from the reference database generally formed separate clusters in the tree (Supplementary Figure 2). As a result it can be expected that when

ASVs are blasted against the references database, the significant matches do not share a common ancestor at the genus level, and as a result the BLCA algorithm cannot assign taxonomy at the genus or species level. In summary, results from this study indicate that, when the reference database is relevant to the sequences amplified (*e.g.* reference AOB *amoA* sequences from marine sediments to assign AOB *amoA* ASVs from marine sediments) the BLCA method is the best approach. On the other hand, if the reference database consists of sequences more dissimilar to the one retrieved from the environment, the NBC method might be more advisable to obtain taxonomic information but the accuracy of this taxonomy might be low. ASVs have substantial merits for the analysis of functional genes for which OTU similarity values are unknown, and in general the field is shifting to ASV based resolution of amplicons e.g. Marshall *et al* (13). Yet, the reference databases used to taxonomically place the amplicons are still based on the traditional threshold based clustering and should improve as more and more ASV based studies become available in the future.

Currently there is a paradigm shift in microbial ecology from OTU to ASV, and this approach is now widely adopted for *16S rRNA* gene studies (29, 32, 93). The use of ASV frees us from using similarity thresholds and produces relevant sequences that can directly be compared between studies; we suggest it offers a pragmatic approach for standardisation of functional gene amplicon sequencing datasets. However, Schloss (94) recently showed that the use of ASV for the *16S rRNA* gene can artificially split bacterial genomes as copies of the gene in a single genome typically do not share 100% similarity. Several functional genes are also present in more than one copy in a genome and their sequences can be slightly different (95, 96). In order to reduce the risk of artificially separating sequences that are in fact identical, we propose to merge together ASVs that translate to the same protein. This allows for the reduction in redundancy in abundance tables by retaining only differences in the gene sequence that result in meaningful biological change (*i.e.* differences in the amino acid sequence of the enzyme). However, whether the clustering at protein level should be done using a threshold lower than 100% is still to be understood among the array of different functional genes and will require genome level resolution, and understanding of active sites on the protein.

## Conclusion

Besides the obvious advantage of not relying on an arbitrary threshold, the ASVs method reflects the sequence diversity of any given functional gene in the environment. This, alongside the ability to compare ASVs among different studies, makes ASV more practical than OTUs for functional gene analysis in environmental microbiology. We also propose that the resulting ASVs should be merged together if they translate to the same protein. Finally, we recommend the use of a relevant database that closely resembles the expected sequences from the environment studied along with the BLCA method for taxonomic classification. If such a database cannot be obtained, the NBC approach might yield better results.

## Acknowledgements

We acknowledge the Cytometry and Imaging facilities and the Molecular core facilities in LIENSs laboratory, University of La Rochelle. We acknowledge the following funders: Thomas Crawford Hayes NUIG (awarded to Cindy J Smith [CJS] & Agatha Lisik [AL]), Harry Smith Vacation Studentships (awarded to AL); Mobility grant from the French Embassy in Ireland (awarded to Hélène Agogué [HA] and CJS)); Research Visit Grant; Microbiology Society and Research Student Mobility Grant; University of Glasgow (awarded to Fabien Cholet [FC]) La Rochelle University; Environmental Protection Agency STRIVE Doctoral Scholarship Scheme (2012-W-PhD-6) (awarded to CJS); FC is supported by a University of Glasgow, Engineering-Scottish Water Research Chair (RCSRF1718643) and EPSRC award EP/V030515/1.

## Supplementary

**Supplementary Figure 1: AOB *amoA*, AOA *amoA* and *nrfA* phylogenetic trees with OTUs and ASVs.** For each gene, phylogenetic trees were drawn with all OTUs/ASVs (top) or with nodes leading only to OTUs/ASVs collapsed (bottom)

**Supplementary Figure 2.** Phylogenetic relationship between the to 50 *nrfA* ASVs and sequences from the referencedatabase covering 125 different genus.

## References

1. Shokralla S, Spall JL, Gibson JF, Hajibabaei M. 2012. Next-generation sequencing technologies for environmental DNA research. Mol Ecol 21:1794– 1805.

2. Shendure J, Balasubramanian S, Church GM, Gilbert W, Rogers J, Schloss JA, Waterston RH. 2017. DNA sequencing at 40: Past, present and future. Nature 550.

3. Roux S, Enault F, le Bronner G, Debroas D. 2011. Comparison of 16S rRNA and protein-coding genes as molecular markers for assessing microbial diversity (Bacteria and Archaea) in ecosystems. FEMS Microbiol Ecol 78:617–628.

4. Poirier S, Rué O, Peguilhan R, Coeuret G, Zagorec M, Champomier-Vergès MC, Loux V, Chaillou S. 2018. Deciphering intra-species bacterial diversity of meat and seafood spoilage microbiota using gyrB amplicon sequencing: A comparative analysis with 16S rDNA V3-V4 amplicon sequencing. PLoS One 13:1–26.

5. Sun DL, Jiang X, Wu QL, Zhou NY. 2013. Intragenomic heterogeneity of 16S rRNA genes causes overestimation of prokaryotic diversity. Appl Environ Microbiol 79:5962–5969.

6. Wu YC, Lu L, Wang BZ, Lin XG, Zhu JG, Cai ZC, Yan XY, Jia ZJ. 2011. Long-Term Field Fertilization Significantly Alters Community Structure of Ammonia-Oxidizing Bacteria rather than Archaea in a Paddy Soil. Soil Sci Soc Am J 75:1431–1439.

7. Duff AM, Zhang LM, Smith CJ. 2017. Small-scale variation of ammonia oxidisers within intertidal sediments dominated by ammonia-oxidising bacteria Nitrosomonas sp. amoA genes and transcripts. Sci Rep 7:1–13.

8. Zhang LM, Duff AM, Smith CJ. 2018. Community and functional shifts in ammonia oxidizers across terrestrial and marine (soil/sediment) boundaries in two coastal Bay ecosystems. Environ Microbiol 20:2834–2853.

9. Aigle A, Prosser JI, Gubry-Rangin C. 2019. The application of high-throughput sequencing technology to analysis of amoA phylogeny and environmental niche specialisation of terrestrial bacterial ammonia-oxidisers. Environ Microbiome 14:3.

10. Cholet F, Ijaz UZ, Smith CJ. 2019. Differential ratio amplicons (R amp) for the evaluation of RNA integrity extracted from complex environmental samples. Environ Microbiol 21:827–844.

11. Cholet F, Ijaz UZ, Smith CJ. 2020. Reverse transcriptase enzyme and priming strategy affect quantification and diversity of environmental transcripts. Environ Microbiol 22:2383–2402.

12. Zhang Y, Wang X, Zhen Y, Mi T, He H, Yu Z. 2017. Microbial diversity and community structure of sulfate-reducing and sulfur-oxidizing bacteria in sediment cores from the East China Sea. Front Microbiol 8:1–17.

13. Marshall IPG, Ren G, Jaussi M, Lomstein BA, Jørgensen BB, Røy H, Kjeldsen KU. 2019. Environmental filtering determines family-level structure of sulfate-reducing microbial communities in subsurface marine sediments. ISME J 13:1920–1932.

14. Dziewit L, Pyzik A, Romaniuk K, Sobczak A, Szczesny P, Lipinski L, Bartosik D, Drewniak L. 2015. Novel molecular markers for the detection of methanogens and phylogenetic analyses of methanogenic communities. Front Microbiol 6:1–12.

15. Varjani SJ, Upasani VN. 2017. A new look on factors affecting microbial degradation of petroleum hydrocarbon pollutants. Int Biodeterior Biodegrad 120:71–83.

16. Liu LY, Xie GJ, Xing DF, Liu BF, Ding J, Cao GL, Ren NQ. 2021. Sulfate dependent ammonium oxidation: A microbial process linked nitrogen with sulfur cycle and potential application. Environ Res 192:110282.

17. Fournier PE, Dubourg G, Raoult D. 2014. Clinical detection and characterization of bacterial pathogens in the genomics era. Genome Med 6:1– 15.

18. Edgar RC. 2018. Updating the 97% identity threshold for 16S ribosomal RNA OTUs. Bioinformatics 34:2371–2375.

19. Green J, Bohannan BJM. 2006. Spatial scaling of microbial biodiversity. Trends Ecol Evol 21:501–507.

20. Horner-Devine MC, Lage M, Hughes JB, Bohannan BJM. 2004. A taxa-area relationship for bacteria. Nature 432:750–753.

21. Storch D, Šizling AL. 2008. The concept of taxon invariance in ecology: Do diversity patterns vary with changes in taxonomic resolution? Folia Geobot 43:329–344.

22. Chen W, Zhang CK, Cheng Y, Zhang S, Zhao H. 2013. A Comparison of Methods for Clustering 16S rRNA Sequences into OTUs. PLoS One 8.

23. Brown EA, Chain FJJ, Crease TJ, Macisaac HJ, Cristescu ME. 2015. Divergence thresholds and divergent biodiversity estimates: Can metabarcoding reliably describe zooplankton communities? Ecol Evol 5:2234– 2251.

24. Koeppel AF, Wu M. 2013. Surprisingly extensive mixed phylogenetic and ecological signals among bacterial Operational Taxonomic Units. Nucleic Acids Res 41:5175–5188.

25. Schmidt TSB, Matias Rodrigues JF, von Mering C. 2015. Limits to robustness and reproducibility in the demarcation of operational taxonomic units. Environ Microbiol 17:1689–1706.

26. Callahan BJ, McMurdie PJ, Holmes SP. 2017. Exact sequence variants should replace operational taxonomic units in marker-gene data analysis. ISME J 11:2639–2643.

27. Caruso V, Song X, Asquith M, Karstens L. 2019. Performance of Microbiome Sequence Inference Methods in Environments with Varying Biomass. mSystems 4:1–19.

28. Moossavi S, Atakora F, Fehr K, Khafipour E. 2020. Biological observations in microbiota analysis are robust to the choice of 16S rRNA gene sequencing processing algorithm: Case study on human milk microbiota. BMC Microbiol 20:1–9.

29. Pauvert C, Buée M, Laval V, Edel-Hermann V, Fauchery L, Gautier A, Lesur I, Vallance J, Vacher C. 2019. Bioinformatics matters: The accuracy of plant and soil fungal community data is highly dependent on the metabarcoding pipeline. Fungal Ecol 41:23–33.

30. Prodan A, Tremaroli V, Brolin H, Zwinderman AH, Nieuwdorp M, Levin E. 2020. Comparing bioinformatic pipelines for microbial 16S rRNA amplicon sequencing. PLoS One 15:1–19.

31. Glassman SI, Martiny JBH. 2018. Broadscale Ecological Patterns Are Robust to Use of Exact Sequence Variants versus Operational Taxonomic Units. mSphere 3:e00148–18.

32. Joos L, Beirinckx S, Haegeman A, Debode J, Vandecasteele B, Baeyen S, Goormachtig S, Clement L, De Tender C. 2020. Daring to be differential: metabarcoding analysis of soil and plant-related microbial communities using amplicon sequence variants and operational taxonomical units. BMC Genomics 21:1–17.

33. Wang Q, Garrity GM, Tiedje JM, Cole JR. 2007. Naïve Bayesian classifier for rapid assignment of rRNA sequences into the new bacterial taxonomy. Appl Environ Microbiol 73:5261–5267.

34. Vinje H, Liland KH, Almøy T, Snipen L. 2015. Comparing K-mer based methods for improved classification of 16S sequences. BMC Bioinformatics 16:1–13.

35. Gao X, Lin H, Revanna K, Dong Q. 2017. A Bayesian taxonomic classification method for 16S rRNA gene sequences with improved species-level accuracy. BMC Bioinformatics 18:1–10.

36. Thom C, Smith CJ, Moore G, Weir P, Ijaz UZ. 2022. Microbiomes in drinking water treatment and distribution: a meta-analysis from source to tap. Water Res 118106.

37. Laima M, Brossard D, Girard M, Richard P, Gouleau D, Joassard L. 2002. The influence of long emersion on biota, ammonium fluxes and nitrification in intertidal sediments of Marennes-Oléron Bay, France. Mar Environ Res 53:381–402.

38. Laima MJC, Girard MF, Vouvé F, Richard P, Blanchard G, Gouleau D. 1999. Nitrification rates related to sedimentary structures in an Atlantic intertidal mudflat, Marennes-Oleron Bay, France. Mar Ecol Prog Ser 191:33–41.

39. Liu Y, Zhou H, Wang J, Liu X, Cheng K, Li L, Zheng J, Zhang X, Zheng J, Pan G. 2015. Short-term response of nitrifier communities and potential nitrification activity to elevated CO2 and temperature interaction in a Chinese paddy field. Appl Soil Ecol 96:88–98.

40. Fang Y, Wang F, Jia X, Chen J. 2019. Distinct responses of ammonia-oxidizing bacteria and archaea to green manure combined with reduced chemical fertilizer in a paddy soil. J Soils Sediments 19:1613–1623.

41. Nair RR, Boobal R, Vrinda S, Bright Singh IS, Valsamma J. 2019. Ammonia-oxidizing bacterial and archaeal communities in tropical bioaugmented zero water exchange shrimp production systems. J Soils Sediments 19:2126–2142.

42. Baskaran V, Patil PK, Antony ML, Avunje S, Nagaraju VT, Ghate SD, Nathamuni S, Dineshkumar N, Alavandi S V., Vijayan KK. 2020. Microbial community profiling of ammonia and nitrite oxidizing bacterial enrichments from brackishwater ecosystems for mitigating nitrogen species. Sci Rep 10:1– 11.

43. Xia W, Zhang C, Zeng X, Feng Y, Weng J, Lin X, Zhu J, Xiong Z, Xu J, Cai Z, Jia Z. 2011. Autotrophic growth of nitrifying community in an agricultural soil. ISME J 5:1226–1236.

44. Mosier AC, Francis CA. 2008. Relative abundance and diversity of ammonia-oxidizing archaea and bacteria in the San Francisco Bay estuary. Environ Microbiol 10:3002–3016.

45. Peng X, Yando E, Hildebrand E, Dwyer C, Kearney A, Waciega A, Valiela I, Bernhard AE. 2012. Differential responses of ammonia-oxidizing archaea and bacteria to long-term fertilization in a New England salt marsh. Front Microbiol 3:1–11.

46. Smith JM, Mosier AC, Francis CA. 2014. Spatiotemporal Relationships Between the Abundance, Distribution, and Potential Activities of Ammonia-Oxidizing and Denitrifying Microorganisms in Intertidal Sediments. Microb Ecol 69:13–24.

47. Sun R, Myrold DD, Wang D, Guo X, Chu H. 2019. AOA and AOB communities respond differently to changes of soil pH under long-term fertilization. Soil Ecol Lett 1:126–135.

48. Abell GCJ, Banks J, Ross DJ, Keane JP, Robert SS, Revill AT, Volkman JK. 2011. Effects of estuarine sediment hypoxia on nitrogen fluxes and ammonia oxidizer gene transcription. FEMS Microbiol Ecol 75:111–122.

49. Baolan H, Shuai L, Lidong S, Ping Z, Xiangyang X, Liping L. 2012. Effect of Different Ammonia Concentrations on Community Succession of Ammonia-oxidizing Microorganisms in a Simulated Paddy Soil Column. PLoS One 7.

50. Pester M, Rattei T, Flechl S, Gröngröft A, Richter A, Overmann J, Reinhold-Hurek B, Loy A, Wagner M. 2012. AmoA-based consensus phylogeny of ammonia-oxidizing archaea and deep sequencing of amoA genes from soils of four different geographic regions. Environ Microbiol 14:525–539.

51. Stempfhuber B, Richter-Heitmann T, Regan KM, Kölbl A, Wüst PK, Marhan S, Sikorski J, Overmann J, Friedrich MW, Kandeler E, Schloter M. 2016. Spatial interaction of archaeal ammonia-oxidizers and nitrite-oxidizing bacteria in an unfertilized grassland soil. Front Microbiol 6:1–15.

52. Biller SJ, Mosier AC, Wells GF, Francis CA. 2012. Global biodiversity of aquatic ammonia-oxidizing archaea: Is partitioned by habitat. Front Microbiol 3:1–15.

53. Lund MB, Smith JM, Francis C a. 2012. Diversity, abundance and expression of nitrite reductase (nirK)-like genes in marine thaumarchaea. ISME J 6:1966– 1977.

54. Wei W, Isobe K, Nishizawa T, Zhu L, Shiratori Y, Ohte N, Koba K, Otsuka S, Senoo K. 2015. Higher diversity and abundance of denitrifying microorganisms in environments than considered previously. ISME J 9:1954– 1965.

55. Shi R, Xu S, Qi Z, Huang H, Liang Q. 2019. Seasonal patterns and environmental drivers of nirS- and nirK-encoding denitrifiers in sediments of Daya Bay, China. Oceanologia 61:308–320.

56. Aalto SL, Saarenheimo J, Arvola L, Tiirola M, Huotari J, Rissanen AJ. 2019. Denitrifying microbial communities along a boreal stream with varying land-use. Aquat Sci 81:1–10.

57. Thompson KA, Bent E, Abalos D, Wagner-Riddle C, Dunfield KE. 2016. Soil microbial communities as potential regulators of in situ N2O fluxes in annual and perennial cropping systems. Soil Biol Biochem 103:262–273.

58. Papaspyrou S, Smith CJ, Dong LF, Whitby C, Dumbrell AJ, Nedwell DB. 2014. Nitrate reduction functional genes and nitrate reduction potentials persist in deeper estuarine sediments why? PLoS One 9.

59. Bu C, Wang Y, Ge C, Ahmad HA, Gao B, Ni SQ. 2017. Dissimilatory Nitrate Reduction to Ammonium in the Yellow River Estuary: Rates, Abundance, and Community Diversity. Sci Rep 7:1–11.

60. Ramanathan B, Boddicker AM, Roane TM, Mosier AC. 2017. Nitrifier gene abundance and diversity in sediments impacted by acid mine drainage. Front Microbiol 8:1–16.

61. Agogué H, Mallet C, Orvain F, De Crignis M, Mornet F, Dupuy C. 2014. Bacterial dynamics in a microphytobenthic biofilm: A tidal mesocosm approach. J Sea Res 92:36–45.

62. Lavergne C, Agogué H, Leynaert A, Raimonet M, De Wit R, Pineau P, Bréret M, Lachaussée N, Dupuy C. 2017. Factors influencing prokaryotes in an intertidal mudflat and the resulting depth gradients. Estuar Coast Shelf Sci 189:74–83.

63. Griffiths RI, Whiteley AS, O’Donnell AG, Bailey MJ. 2000. Rapid method for coextraction of DNA and RNA from natural environments for analysis of ribosomal DNA- and rRNA-based microbial community composition. Appl Environ Microbiol 66:5488–5491.

64. Hornek R, Pommerening-Röser A, Koops HP, Farnleitner AH, Kreuzinger N, Kirschner A, Mach RL. 2006. Primers containing universal bases reduce multiple amoA gene specific DGGE band patterns when analysing the diversity of beta-ammonia oxidizers in the environment. J Microbiol Methods 66:147– 155.

65. Wuchter C, Abbas B, Coolen MJL, Herfort L, van Bleijswijk J, Timmers P, Strous M, Teira E, Herndl GJ, Middelburg JJ, Schouten S, Sinninghe Damsté JS. 2006. Archaeal nitrification in the ocean. Proc Natl Acad Sci U S A 103:12317–22.

66. Hallin S, Lindgren PE. 1999. PCR detection of genes encoding nitrite reductase in denitrifying bacteria. Appl Environ Microbiol 65:1652–1657.

67. Levy-Booth DJ, Prescott CE, Grayston SJ. 2014. Microbial functional genes involved in nitrogen fixation, nitrification and denitrification in forest ecosystems. Soil Biol Biochem.

68. Welsh A, Chee-Sanford JC, Connor LM, Löffler FE, Sanford RA. 2014. Refined NrfA phylogeny improves PCR-based nrfA gene detection. Appl Environ Microbiol 80:2110–2119.

69. Pester M, Maixner F, Berry D, Rattei T, Koch H, Lücker S, Nowka B, Richter A, Spieck E, Lebedeva E, Loy A, Wagner M, Daims H. 2014. NxrB encoding the beta subunit of nitrite oxidoreductase as functional and phylogenetic marker for nitrite-oxidizing Nitrospira. Environ Microbiol 16:3055–3071.

70. Sickle NAJ, Fass. JN. 2011. A sliding-window, adaptive, quality-based trimming tool for FastQ file 1–9.

71. Schirmer M, Ijaz UZ, D’Amore R, Hall N, Sloan WT, Quince C. 2015. Insight into biases and sequencing errors for amplicon sequencing with the Illumina MiSeq platform. Nucleic Acids Res 43.

72. D’Amore R, Ijaz UZ, Schirmer M, Kenny JG, Gregory R, Darby AC, Shakya M, Podar M, Quince C, Hall N. 2016. A comprehensive benchmarking study of protocols and sequencing platforms for 16S rRNA community profiling. BMC Genomics 17.

73. Callahan BJ, McMurdie PJ, Rosen MJ, Han AW, Johnson AJA, Holmes SP. 2016. DADA2: High-resolution sample inference from Illumina amplicon data. Nat Methods 13:581–583.

74. Katoh K, Asimenos G, Toh H. 2009. Multiple Alignment of DNA Sequences with MAFFT, p. 39–64. *In* Bioinformatics for DNA Sequence Analysis, Methods in Molecular Biology.

75. Price MN, Dehal PS, Arkin AP. 2010. FastTree 2 - Approximately maximum-likelihood trees for large alignments. PLoS One 5.

76. Winter DJ. 2017. rentrez: An R package for the NCBI eUtils API. R J 9:520– 526.

77. Caporaso JG, Kuczynski J, Stombaugh J, Bittinger K, Bushman FD, Costello EK, Fierer N, Peña AG, Goodrich JK, Gordon JI, Huttley G a, Kelley ST, Knights D, Koenig JE, Ley RE, Lozupone C a, Mcdonald D, Muegge BD, Pirrung M, Reeder J, Sevinsky JR, Turnbaugh PJ, Walters W a, Widmann J, Yatsunenko T, Zaneveld J, Knight R. 2010. QIIME allows analysis of high-throughput community sequencing data. Nat Methods 7:335–336.

78. Charif D, Lobry JR. 2007. SeqinR 1.0-2: a contributed package to the R project for statistical computing devoted to biological sequences retrieval and analysis Delphine, p. 207–232. *In* Structural approaches to sequence evolution: Molecules, networks, populations.

79. Kumar S, Stecher G, Tamura K. 2016. MEGA7: Molecular Evolutionary Genetics Analysis Version 7.0 for Bigger Datasets. Mol Biol Evol 33:1870– 1874.

80. Yu G, Smith DK, Zhu H, Guan Y, Lam TTY. 2017. Ggtree: an R Package for Visualization and Annotation of Phylogenetic Trees With Their Covariates and Other Associated Data. Methods Ecol Evol 8:28–36.

81. Prosser JI, Nicol GW. 2012. Archaeal and bacterial ammonia-oxidisers in soil: The quest for niche specialisation and differentiation. Trends Microbiol. Elsevier Ltd.

82. Ke X, Angel R, Lu Y, Conrad R. 2013. Niche differentiation of ammonia oxidizers and nitrite oxidizers in rice paddy soil. Environ Microbiol 15:2275– 2292.

83. Shen JP, Xu Z, He JZ. 2014. Frontiers in the microbial processes of ammonia oxidation in soils and sediments. J Soils Sediments.

84. Gao D, Liu F, Xie Y, Liang H. 2018. Temporal and spatial distribution of ammonia-oxidizing organisms of two types of wetlands in Northeast China. Appl Microbiol Biotechnol 102:7195–7205.

85. Hink L, Gubry-Rangin C, Nicol GW, Prosser JI. 2018. The consequences of niche and physiological differentiation of archaeal and bacterial ammonia oxidisers for nitrous oxide emissions. ISME J 12:1084–1093.

86. Hou L, Xie X, Wan X, Kao S-J, Jiao N, Zhang Y. 2018. Niche differentiation of ammonia and nitrite oxidizers along a salinity gradient from the Pearl River estuary to the South China Sea. Biogeosciences Discuss 1–61.

87. Aigle A, Gubry-Rangin C, Thion C, Estera-Molina KY, Richmond H, Pett-Ridge J, Firestone MK, Nicol GW, Prosser JI. 2020. Experimental testing of hypotheses for temperature- and pH-based niche specialization of ammonia oxidizing archaea and bacteria. Environ Microbiol 22:4032–4045.

88. Jia Z, Zhou X, Xia W, Fornara D, Wang B, Wasson EA, Christie P, Polz MF, Myrold DD. 2020. Evidence for niche differentiation of nitrifying communities in grassland soils after 44 years of different field fertilization scenarios. Pedosphere 30:87–97.

89. Schloss PD. 2010. The effects of alignment quality, distance calculation method, sequence filtering, and region on the analysis of 16S rRNA gene-based studies. PLoS Comput Biol 6:19.

90. Franzén O, Hu J, Bao X, Itzkowitz SH, Peter I, Bashir A. 2015. Improved OTU-picking using long-read 16S rRNA gene amplicon sequencing and generic hierarchical clustering. Microbiome 3:43.

91. García-López R, Cornejo-granados F, Lopez-zavala AA, Cota-Huízar A, Sotelo-Mundo RR, Gómez-Gil B, Ochoa-Leyva A. 2021. OTUs and ASVs Produce Comparable Taxonomic and Diversity using tailored abundance filters. Genes (Basel) 12:564.

92. Jones CM, Stres B, Rosenquist M, Hallin S. 2008. Phylogenetic analysis of nitrite, nitric oxide, and nitrous oxide respiratory enzymes reveal a complex evolutionary history for denitrification. Mol Biol Evol 25:1955–1966.

93. García-García N, Tamames J, Linz AM, Pedrós-Alió C, Puente-Sánchez F. 2019. Microdiversity ensures the maintenance of functional microbial communities under changing environmental conditions. ISME J 13:2969–2983.

94. Schloss PD. 2021. Amplicon sequence variants artificially split bacterial genomes into separate clusters. bioRxiv 1–12.

95. Chain P, Lamerdin J, Larimer F, Regala W, Lao V, Land M, Hauser L, Alan H, Klotz M, Norton J, Sayavedra-Soto L, Arciero D, Hommes N, Whittaker M, and Arp D. 2003. Complete Genome Sequence of the Ammonia-Oxidizing Bacterium and Obligate Chemolithoautotroph Nitrosomonas europaea. J Bacteriol 185:2759–2773.

96. Norton JM, Klotz MG, Stein LY, Arp DJ, Bottomley PJ, Chain PSG, Hauser LJ, Land ML, Larimer FW, Shin MW, Starkenburg SR. 2008. Complete genome sequence of Nitrosospira multiformis, an ammonia-oxidizing bacterium from the soil environment. Appl Environ Microbiol 74:3559–3572.

97. Oksanen J, Kindt R, O’Hara RB. 2005. vegan: Community Ecology Package. Available from http://cc.oulu.fi/~jarioksa/3.

98. Hsieh TC, Ma KH, Chao A. 2016. iNEXT: an R package for rarefaction and extrapolation of species diversity (Hill numbers). Methods Ecol Evol 7:1451– 1456.

99. Legendre P, Gallagher ED. 2001. Ecologically meaningful transformations for ordination of species data. Oecologia 129:271–280.

